# Predictions for the binding domain and potential new drug targets of 2019-nCoV

**DOI:** 10.1101/2020.02.26.961938

**Authors:** Zehua Zeng, Zhi Luo, Hongwu Du

## Abstract

An outbreak of new SARS-like viral in Wuhan, China has been named 2019-nCoV. The current state of the epidemic is increasingly serious, and there has been the urgent necessity to develop an effective new drug. In previous studies, it was found that the conformation change in CTD1 was the region where SARS-CoV bound to human ACE2. Although there are mutations of the 2019-nCoV, the binding energy of ACE2 remains high. The surface glycoprotein of 2019-nCoV was coincident with the CTD1 region of the S-protein by comparing the I-TASSER prediction model with the actual SARS model, which suggests that 2019-nCoV may bind to the ACE2 receptor through conformational changes. Furthermore, site prediction on the surface glycoprotein of 2019-nCoV suggests some core amino acid area may be a novel drug target against 2019-nCoV.

## Introduction

In the past of 18 years, many researchers have done lots of work on etiology mechanism related to SARS, and Spike glycoprotein (S-protein) was confirmed to be a key for entering human cell by binding with angiotensin converting enzyme 2 (ACE2) receptor^1^.

Recently, a new type of pneumonia which is caused by 2019-nCoV has outbroken nationwide in China and the previous studies have shown that 2019-nCoV is like SARS-CoV. In the last few days, several 2019-nCoV strains have been successfully separated from patients and the results of sequencing data can be acquired on Sharing Avian Influenza Data and GenBank.

According to the evolutionary analysis of coronavirus, 2019-nCov is probably originated from bat, and the S-protein of 2019-nCoV may enter human cells by interacting with human ACE2 receptor, which revealed the pathopoeia mechanism of 2019-nCoV^1^. On the other hand, 2019-nCoV shares about 96.2% sequence identity to BatCoV RATG13^2^. By comparing the amino acid sequence of 2019-nCoV S-protein (GenBank Accession: MN908947.3) with Bat SARS-like coronavirus isolate bat-SL-CoVZC45 and Bat SARS-like coronavirus isolate Bat-SL-CoVZXC21, the latter two were shown to share 89.1% and 88.6% sequence identity to 2019- nCoV S-protein (supplementary figure 1). Thus, the hypothesis that 2019-nCoV may share the same pathway with Bat SARS-like coronavirus.

However, the existing studies used swiss-model need further optimization, for example, the ligand binding site on the predicted S-protein structure are still not clear. Additionally, the template only shares 76.4% sequence identity, which means the result may not fully reflect the reality. Thus, a new method to predict the threedimensional structure and related functions of proteins from another perspective and most importantly, a new method to predict potential drug target may need to be applied.

Iterative Threading Assembly Refinement (I-TASSER) is a layered method for protein structure and function prediction^3^. It firstly identifies the structural template from the PDB through the local meta-threaded method (LOMETS), and then constructs the full-length atomic model through the fragment assembly simulation based on the iterative template. Next, the 3D model is re-woven into the thread through BioLiP, the protein function database, to yield a functional insight into the target. In the recent protein structure predictions for CASP7, CASP8, CASP9, CASP10, CASP11, CASP12 and CASP13, I-TASSER remains one of the best method for the automated protein structure prediction^3^.

Phyre2 is a tool which uses remote homology detection methods to build 3D models, predict ligand binding sites and analyse the effect of amino-acid variants. The difference between Phyre2 and other methods is not accuracy but ease of use^4^.

Therefore, I-TASSER and Phyre2 were used to predict the structure of spike protein in Wuhan pneumonia (GenBank: MN908947.3).

## Materials and Methods

### Sequence Data Collection

The latest 2019-ncov sequences (Supplementary Table 1; in total 33 strains analyzed) from GISAID and MN908947.3 sequences published on NCBI were collected and analyzed. MEGA(Version 7.026) was used to align 33 nucleotide sequences. The aligned nucleotide sequences were constructed by NJ method.

### Analysis of Virus Variability

Using online tools blast (https://blast.ncbi.nlm.nih.gov/Blast.cgi) to analyse MN908947.3 amino acid sequence and the 33 2019-nCoV sequences downloaded from GISAID.

### Prediction by I-TASSER Protein Model

The MN908947.3 spike protein sequence was first extracted from GenBank file, the sequence was then submitted to the online tools (https://zhanglab.ccmb.med.umich.edu/I-TASSER/) which uses existing model to predict protein structure. Parameters were not assigned to other constraints, all template libraries of I-TASSER were selected, and secondary structures of specific residues were not specified for protein structure analysis.

This section reports biological annotations of the target protein by COFACTOR and COACH based on the I-TASSER structure prediction. COFACTOR deduces protein functions (ligand-binding sites, EC and GO) by using structure comparison and protein-protein networks. COACH is a meta-server approach that combines multiple function annotation results (on ligand-binding sites) from COFACTOR, TM-SITE and S-SITE programs.

### Prediction by Phyre2 Protein Model

The MN908947.3 spike protein amino acid sequence was submitted to the online tools (http://www.sbg.bio.ic.ac.uk/phyre2/html/page.cgi?id=index) which uses existing model to predict protein sequences.

## Results

No Significant Variation was Found in 2019-nCoV

So far, variability analysis of the known 33 coronavirus sequences showed that during the one-month period of transmission of pneumonia in Wuhan (2019.12.24-2020.01.29), all viral RNA sequences had a Query Cover greater than 99.9% compared with the sequence was matched (MN908947.3). There was no significant variation in the virus, so the analysis of MN908947.3 could be applied to the prevalent Wuhan pneumonia species (excluding BetaCoV/Kanagawa/1/2020|EPI_ISL_402126).

In addition, in the phylogenetic tree, the evolution of 33 virus samples was divided into three categories. Clade A is basically unchanged nucleotide sequence, Clade B is A sample with individual site variation (site<=3), and Clade C is A single sample (site=5). The distance scale of the tree is 0.0001. In the figure 1, we can see that the distance of the sample with the greatest difference (Clade C) is only 0.0002. In summary, no significant variation has occurred in the known virus samples.

**Fig.1.**
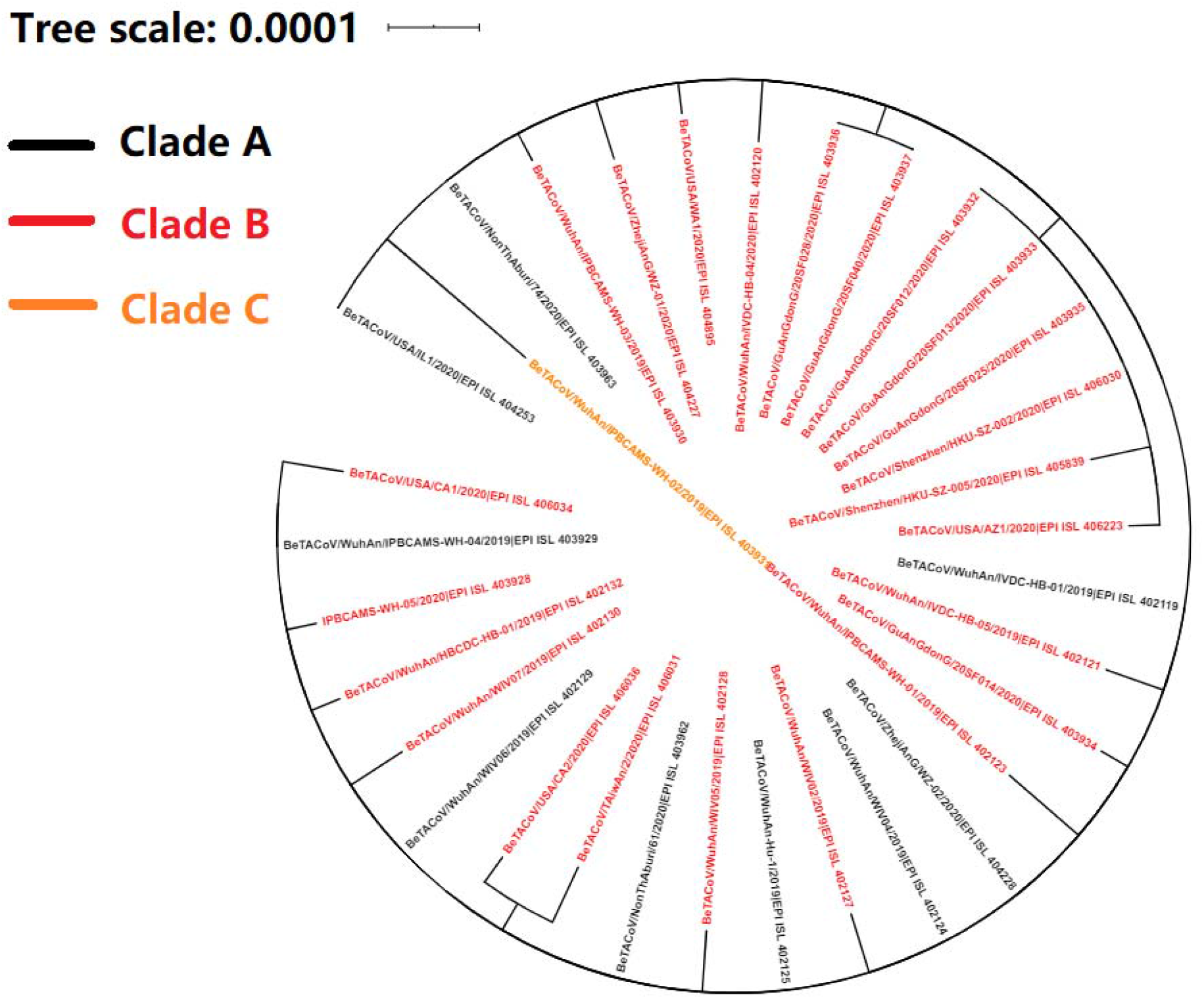
Evolutionary relationships. The evolutionary history was inferred using the Neighbor-Joining method^14^. The optimal tree with the sum of branch length = 0.00123858 is shown. The tree is drawn to scale, with branch lengths in the same units as those of the evolutionary distances used to infer the phylogenetic tree. The evolutionary distances were computed using the Maximum Composite Likelihood method and are in the units of the number of base substitutions per site. The analysis involved 33 nucleotide sequences. Codon positions included were 1st+2nd+3rd+Noncoding. All positions containing gaps and missing data were eliminated. There were a total of 29874 positions in the final dataset. Evolutionary analyses were conducted in MEGA7^15^. Decorate by iTol^16^. Black is Clade A; Red is Clade B; Orange is Clade C. The data up to 2020.1.30.

### Prediction of 2019-nCoV Surface Glycoprotein Structure Model

**I-TASSER:** The results showed that five predicted protein structure models (named as model1-5, respectively) were co-existed, and the changes of normalized B factor^5^ of each model were shown in figure 2. A total of 5 models were generated, which were sorted according to the clustering size, and the local structure accuracy curve of each model was shown (supplementary figure 3). The local accuracy is defined as the distance deviation (in Angstrom) between the position of residues in the model and the original structure, and the size of its C score is given in Supplementary Table 2.

**Fig.2.**
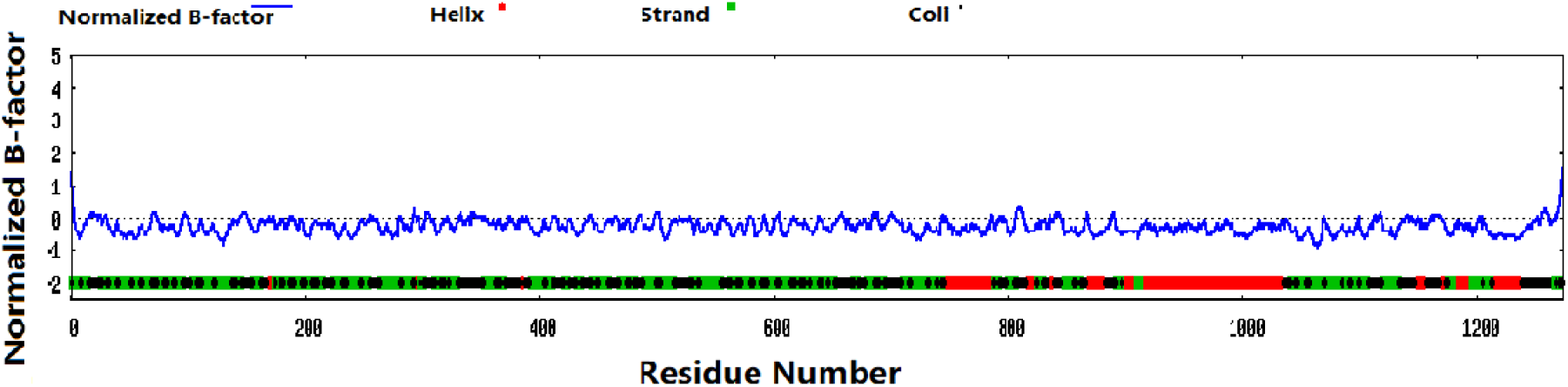
Sequences of 2019-nCoV. The abscissa represents the sequence of nucleotide sequences of 2019-nCoV, and the ordinate represents the magnitude of normalized b-factor values.

**Phyre2:** Phyre2 analytical model is based on c5×5b B (PDB ID) (supplementary figure 4), which models 1049 residues (82.1% of the sequence) with a confidence of 100.0% from a single highest scoring template.

In Supplementary Table 3, the top 10 known proteins in I-TASSER with the highest structural similarity to the prediction model with the highest score were listed (all structures in the PDB library were matched by tm-align). After analyzing the structures of these 10 known proteins, it was found that only the first two (PDB ID: 5×58 A and 6nzk A) were coronavirus proteins. Moreover, 5×58, which ranks the first, is the spike trimer of SARS, which is exactly consistent with our prediction that the nucleotide sequence was inferred to be spike. Reports on 5×58 indicated that spike was divided into four structural domains, among which C-terminal domain 1 (CTD1s) binds ACE2 as receptor binding domain (figure 4c)^6^. After combining 5×58 with I-TASSER prediction model (figure 4d), there were similar up and down proteins. This indicated that the binding of the novel coronavirus (2019-nCoV) to human cells and the conformation of CTD1 may also change from downward to upward, making ACE2 binding to RBD unimpeded.

### Functional Prediction of 2019-nCoV Surface Glycoprotein

In the functional prediction model of proteins, binding sites ranking from 1st to 5th were PDB ID^7^: 3srcA (C Sore=0.05), 1a5tA (C Sore= 0.03), 4dv3B (C Sore= 0.03), 4fmal (C Sore= 0.03), and 5hhja (C Sore= 0.02) (Supplementary Table 4), respectively. Further analysis of the binding site ranking the first showed that it was the binding site of Acyl-homoserine lactone acylase (PvdQ) which catalyzes the deacylation of acylhomoserine lactone (AHL or acyl-HSL), releasing homoserine lactone (HSL) and the corresponding fatty acid (figure 3), and the residues of binding site were Thr-874, Leu-877, and Leu-878 with binding affinity IC50=130 μM (from pdbbind-cn).

**Fig.3.**
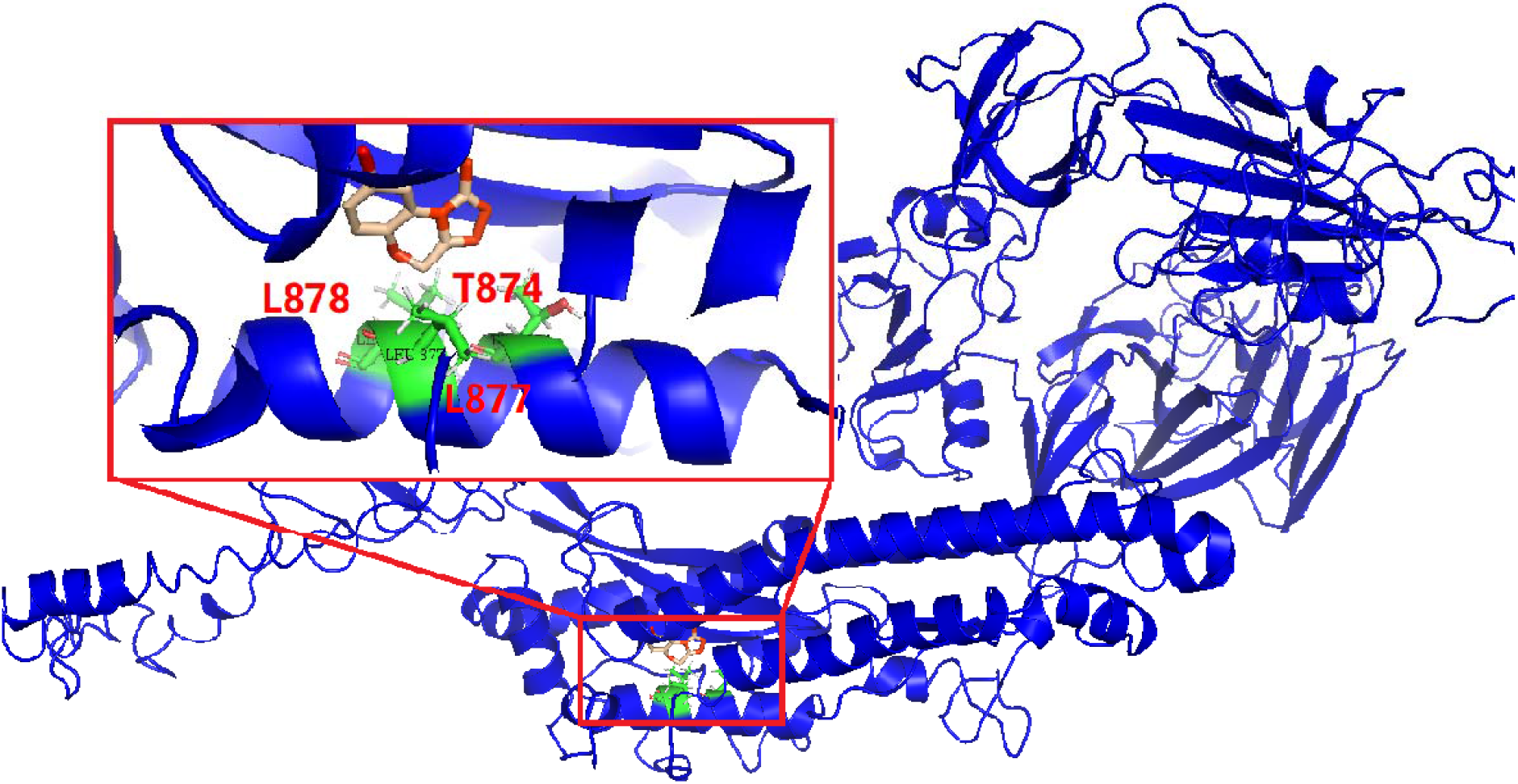
The binding domain. The msata amplifies the protein space construction at the 3srcA sites, the black font represents the residue name, and the blue represents the surface glycoprotein of 2019-nCoV.

### Target For New Drug Development of 2019-nCoV

In the receptor results, 3src was ranked the first (3src is the PDB ID of PvdQ,), and its predicted binding ligand was 28N (8-BROMO-4H-[1,2,4]OXADIAZOLO[3,4-C][1,4]BENZOXAZIN-1-ONE). We hypothesized that the molecules with similar structures like 28N might bind to the site Thr-874, Leu-877, and Leu-878 of the surface glycoprotein of 2019-nCoV, thereby enabling viral hydrolysis and inhibiting viral reproduction.

## Discussion

In this study, the nucleotide interval of the surface glycoprotein of the 2019-nCoV was analyzed and it was similar to the spike interval of SARS-CoV^8^. Three-dimensional structure prediction of protein showed that the prediction model with the highest score in I-TASSER was comparable to the 5×58 spike protein electron microscope results, and the prediction model with the highest score in phyre2 was comparable to the 5×5b SARS-CoV spike protein. Moreover, the prediction model in I-TASSER was highly similar to that in 5×58 which indirectly makes the nucleotide interval 2019-nCoV, indicating that the possibility of glycoprotein expression was increased again.

In addition, the 3D structure of CTD1 in the binding region of SARS-CoV and ACE2 was enlarged. In figure 4a and figure 4d, it was observed that there was a high degree of overlap between the S2 subunit of SARS-CoV and the prediction model of 2019-nCoV, and the prediction model with conformation up and down was in line with the actual model of spike of SARS. Therefore, we speculated that 2019-nCoV and SARS may have the same ACE2 binding model. Previous studies built a preliminary 3D model of 2019-nCoV using the Swiss method, and the researchers found that compared with SARS, the 2019-nCOV changed four key bases, but its 3D conformation did not change significantly^1^. In our 2019-nCoV 3D model, we found that 2019- nCoV was highly similar to the receptor-binding domain (RBD) domain of SARS-CoV. The main chains of the two protein models were highly coincident and the conformation was in the same direction, and the possibility of combining 2019-nCoV with ACE2 was increased from another perspective.

**Fig.4.**
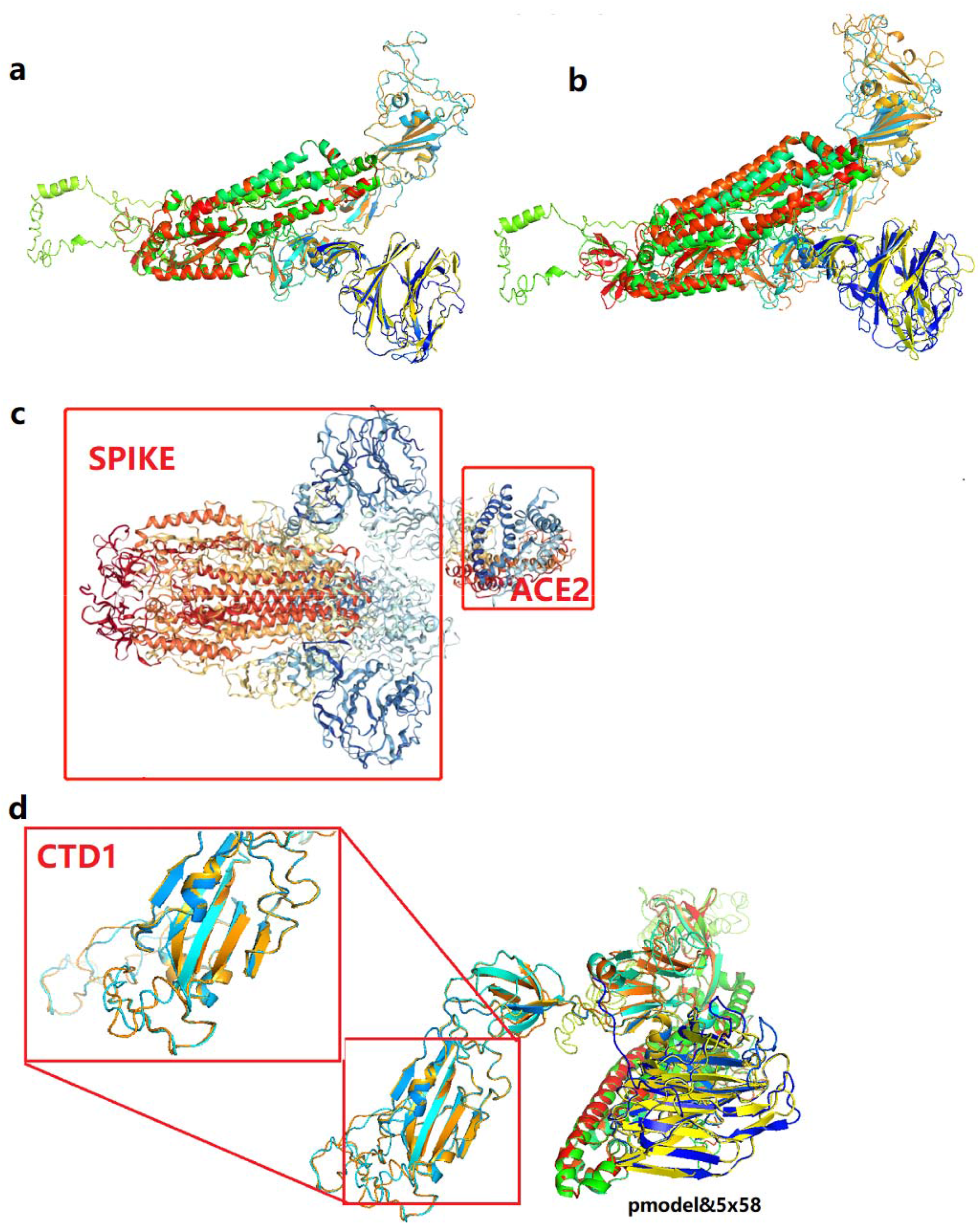
3D structure of CTD1 in the binding region of SARS-CoV and ACE2. **a.** The diagram shows the superposition of glycoprotein structure and 5×58 structure on the surface of the predicted 2019-nCoV, which ranked first. **B.** The diagram shows the overlap between the surface glycoprotein structure and the 5×58 structure of the predicted 2019-nCoV, which ranks second. **C.** The part circled in red represents ace2 and spike trimer in 5×58. **d.** Put 5×58 and the prediction model in the same coordinate system, zoom in on the CTD1 binding region, and the arrow represents the direction of the protein conformation.

In previous reports, ACE2 was highly expressed in alveolar epithelial cells, intestinal epithelial cells^9^, and in the eyes of the aqueous humor, cornea and conjunctiva^10,11^. In the most recent cases, people with conjunctivitis and diarrhea were infected. Therefore, from this perspective, it can also be explained that the entry pathway of 2019-nCov in human may be binding to ACE2 receptor. If this inference is true, then healthy people should also be careful to treat patients’ feces. and people who go to places with high virus density (such as fever clinic) must pay attention to eye protection measures to prevent the viral infection through the Aqueous humor.

Here we suggest that 28N may be a potential drug target for 2019-nCoV. According to Coach analysis, we found that 28N could be an strong ligand to bing the S protein. In past studies, it was found that 28N can strongly bind PvdQ, and the residues near its binding site are T166, L169 and L170^12^. In the spatial structure simulation of S protein of 2019-ncov, we found that S protein has a three-dimensional structure highly cosimilar to PvdQ. More importantly, its binding site residues are also T874, L877 and L878, which are almost exactly matched. This conclusion indicates that 28N can bind to S protein. A search on the pubchem platform revealed that seven of the eight BioAssay Results listed as active were as inhibitors and one as stimulants. We speculate that 28N can inhibit the activity of S protein of 2019-ncov, but more experiments are needed to confirm this conclusion.

Previous reports have suggested that the conformational transformation of CTD1 from bottom to top was a prerequisite for receptor binding^13^, and that the spike protein of 2019-nCoV should also be trimer. A switch with more than one “up” CTD1 conformation will result in cleavage of the S1-ACE2 complex. Therefore, the research on new pneumonia drugs can also start from how to make spike trimer to turn on the upward switch. Antibodies against ACE2 or any compound that can bind to ACE2 are expected to be used in 2019-nCoV therapy, and the recognition of receptor binding sites will also contribute to the development of 2019-nCoV vaccine.

In conclusion, two different deep learning-based models of protein structure prediction were used to show that the binding domain of 2019-nCov was highly conserved to the CTD1 region of the spike protein of SARS. In the functional prediction of protein, 28N or its molecules with similar structuresmay be used as a potential drug target to hydrolyze 2019-nCoV. Hopefully this study can help to develop new drug for the therapy of 2019-nCoV.

## Data and Code availability

All custom user data and code used for the sequence alignment and evolutionary analysis are available from https://github.com/Starlitnightly/Bio_2019_nCoV. The code was written and tested using Python 3.7.

## References

1 Xu, X. et al., Evolution of the novel coronavirus from the ongoing Wuhan outbreak and modeling of its spike protein for risk of human transmission. Science China Life Sciences (2020).

2 Zhou, P. et al., Discovery of a novel coronavirus associated with the recent pneumonia outbreak in humans and its potential bat origin. bioRxiv 2020 (2020).

3 Roy, A., Kucukural, A. & Zhang, Y., I-TASSER: a unified platform for automated protein structure and function prediction. NAT PROTOC 5 725 (2010).

4 Kelley, L. A., Mezulis, S., Yates, C. M., Wass, M. N. & Sternberg, M. J. E., The Phyre2 web portal for protein modeling, prediction and analysis. NAT PROTOC 10 845 (2015).

5 Yang, J., Wang, Y. & Zhang, Y., ResQ: An Approach to Unified Estimation of B-Factor and Residue-Specific Error in Protein Structure Prediction. J MOL BIOL 428 693 (2016).

6 Yuan, Y. et al., Cryo-EM structures of MERS-CoV and SARS-CoV spike glycoproteins reveal the dynamic receptor binding domains. NAT COMMUN 8 15092 (2017).

7 Yang, J., Roy, A. & Zhang, Y., Protein-ligand binding site recognition using complementary binding-specific substructure comparison and sequence profile alignment. BIOINFORMATICS 29 2588 (2013).

8 Ji, W., Wang, W., Zhao, X., Zai, J. & Li, X., Homologous recombination within the spike glycoprotein of the newly identified coronavirus may boost cross-species transmission from snake to human. J MED VIROL (2020).

9 Holappa, M., Vapaatalo, H. & Vaajanen, A., Many Faces of Renin-angiotensin System - Focus on Eye. The Open Ophthalmology Journal 11 122 (2017).

10 Sun, Y., Liu, L., Pan, X. & Jing, M., Mechanism of the action between the SARS-CoV S240 protein and the ACE2 receptor in eyes. INT J OPHTHALMOL-CHI 6 783 (2006).

11 Saitou, N. M. & Nei, M., The neighbor-joining method: a new method for reconstructing phylogenetic trees. MOL BIOL EVOL 24 189 (1987).

12 Drake, E. J. & Gulick, A. M., Structural Characterization and High-Throughput Screening of Inhibitors of PvdQ, an NTN Hydrolase Involved in Pyoverdine Synthesis. ACS CHEM BIOL 6 1277 (2011).

13 Song, W., Gui, M., Wang, X. & Xiang, Y., Cryo-EM structure of the SARS coronavirus spike glycoprotein in complex with its host cell receptor ACE2. PLOS PATHOG 14 e1007236 (2018).

14 Tamura, K., Nei, M. & Kumar, S., Prospects for inferring very large phylogenies by using the neighbor-joining method. Proc Natl Acad Sci U S A 101 11030 (2004).

15 Sudhir, K., Glen, S. & Koichiro, T., MEGA7: Molecular Evolutionary Genetics Analysis Version 7.0 for Bigger Datasets. Molecular Biology & Evolution 7 (2016).

16 Letunic, I. & Bork, P., Interactive Tree Of Life (iTOL) v4: recent updates and new developments. NUCLEIC ACIDS RES 47 W256 (2019).

