## Supplemental Fig.1&2&3 Supplemental Table 1&2&3&4 for "Predictions for the binding domain and potential new drug targets of 2019-nCoV"

**Supplementary**

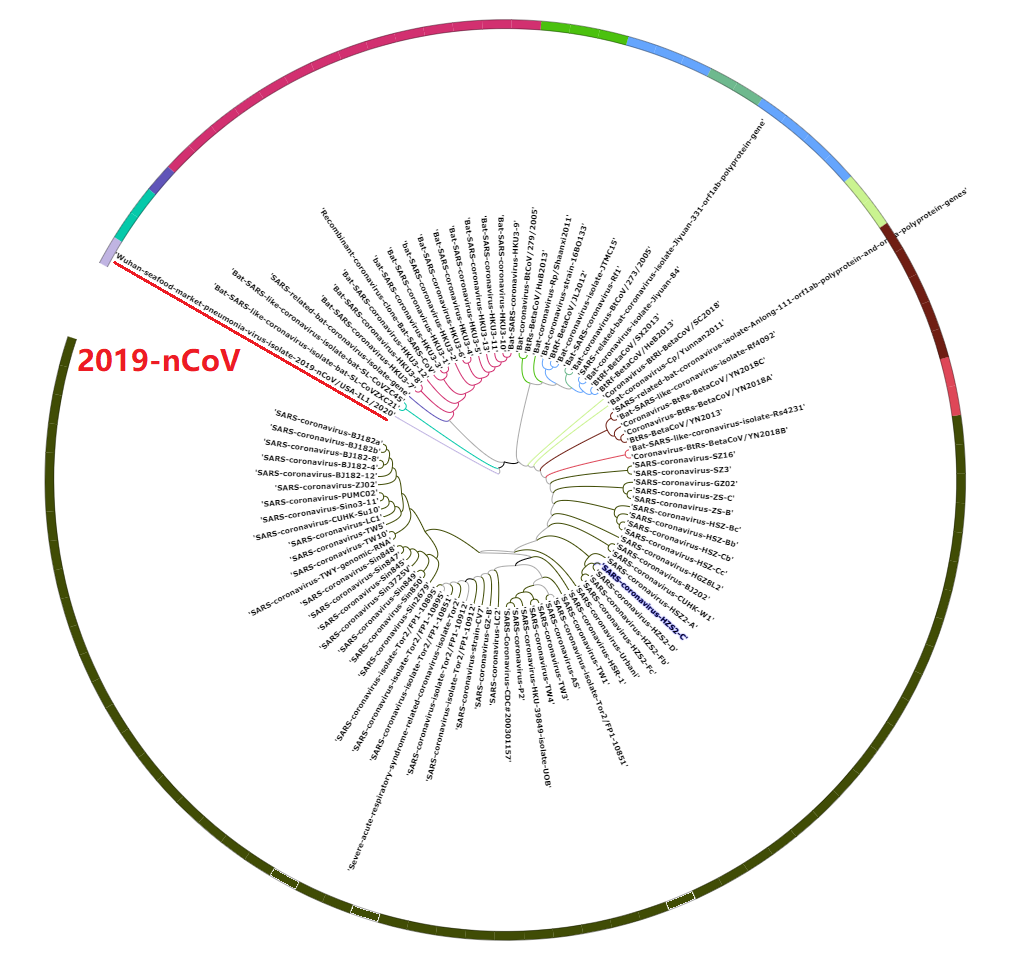

**Supplement Figure 1: System evolution tree.** Marked in Red are the 2019-nCoV nucleotide sequences used for detection, Each color represents a category represent the nucleotide sequences in different species. This figure is described by iTol.

**Supplement Figure 2: The estimated accuracy of every model by I-TASSER.** The abscissa represents the Residue Number, and the ordinate represents Estimated Accuracy. Figure a-e shows the local structure accuracy of each of the five models ranked from 1 to 5.

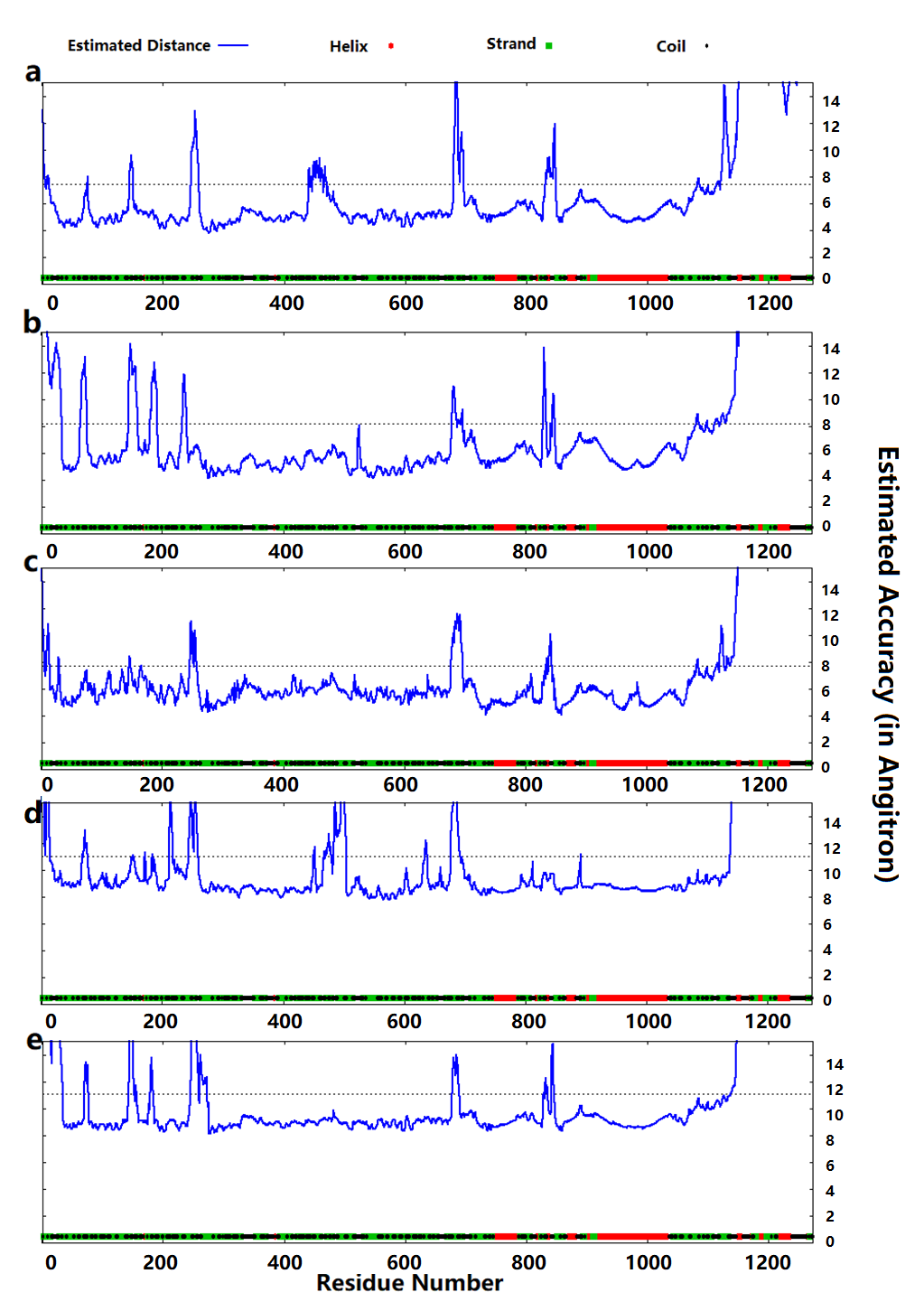

**Supplement Figure 3:** Five structure analyzed. a-e is the predicted glycoprotein structure on the surface of 2019-nCoV, image coloured by rainbow N → C terminus.

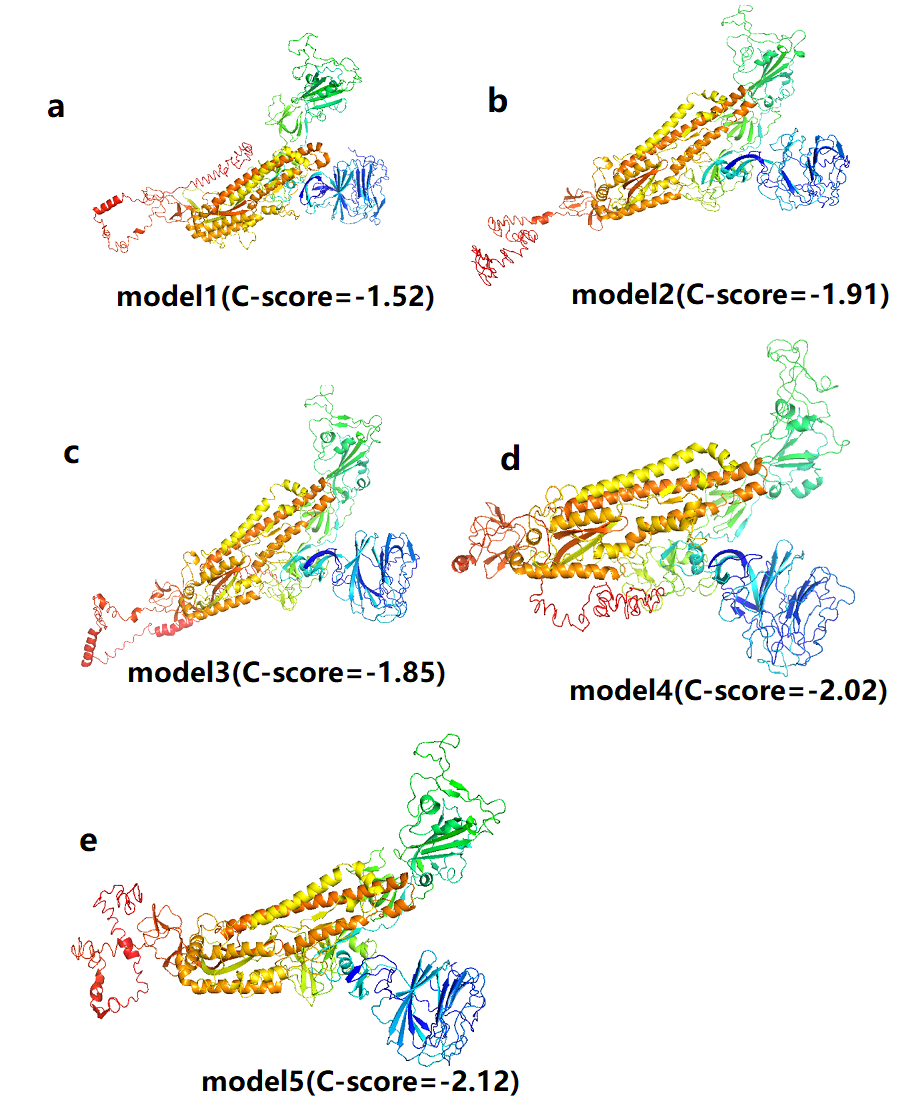

**Supplement Figure 4: a** Phyre2 prediction model, **b** 5x58 actual structure. The region in red is the CTD1 region of the surface glycoprotein.

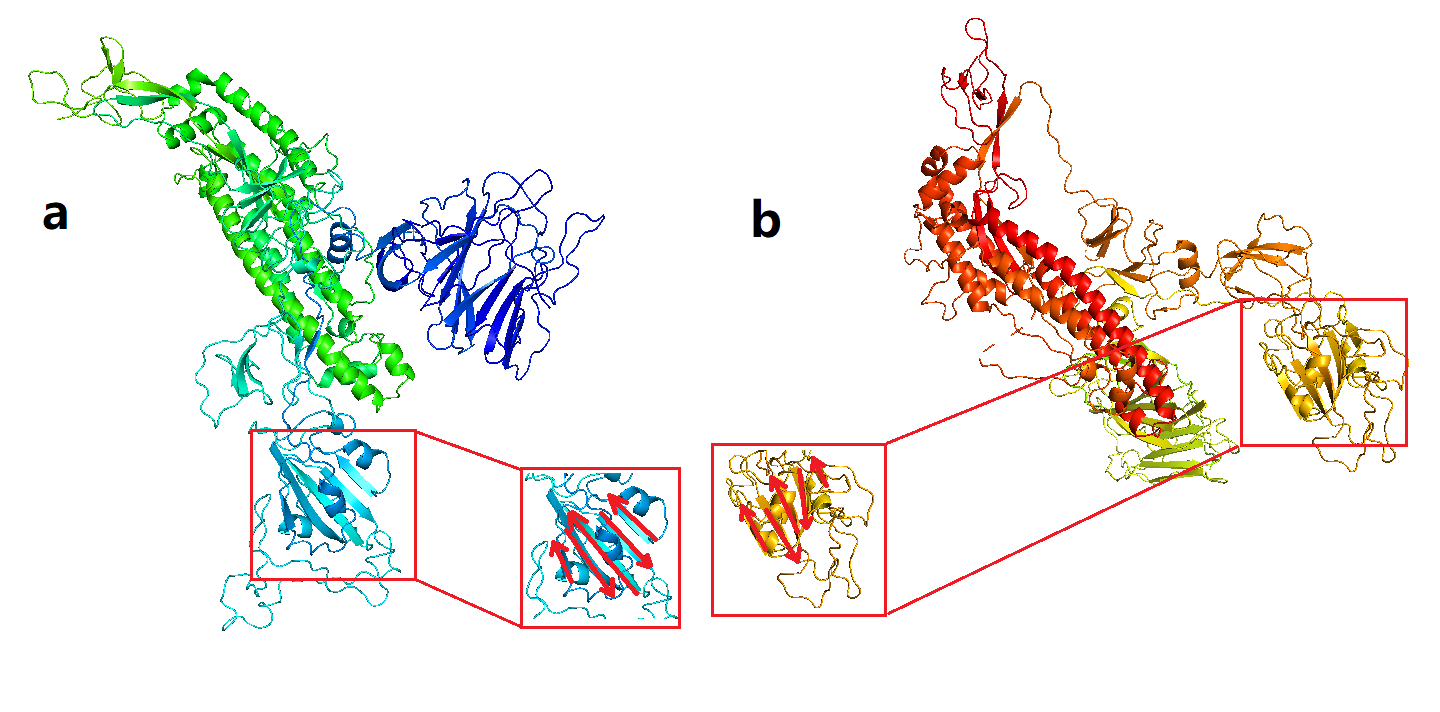

**Supplement Table 1: The blast result of the GISAID sequence and MN908947.3**

| **Description** | **Max Score** | **Total Score** | **Query Cover** | **E value** | **Per.Ident** | **Accession** |
| --- | --- | --- | --- | --- | --- | --- |
| BeTACoV/WuhAn/IVDC-HB-01/2019\|EPI_ISL_402119 | 55182 | 55182 | 99% | 0.0 | 100.00% | Query_26999 |
| BeTACoV/WuhAn/IVDC-HB-04/2020\|EPI_ISL_402120 | 55171 | 55171 | 99% | 0.0 | 99.99% | Query_24673 |
| BeTACoV/WuhAn/IVDC-HB-05/2019\|EPI_ISL_402121 | 55171 | 55171 | 99% | 0.0 | 99.99% | Query_59293 |
| BeTACoV/WuhAn/IPBCAMS-WH-01/2019\|EPI_ISL_402123 | 55166 | 55166 | 99% | 0.0 | 99.99% | Query_59047 |
| BeTACoV/WuhAn/WIV04/2019\|EPI_ISL_402124 | 55182 | 55182 | 99% | 0.0 | 100.00% | Query_64871 |
| BeTACoV/WuhAn-Hu-1/2019\|EPI_ISL_402125 | 55182 | 55182 | 99% | 0.0 | 100.00% | Query_51587 |
| BeTACoV/WuhAn/WIV02/2019\|EPI_ISL_402127 | 55171 | 55171 | 99% | 0.0 | 99.99% | Query_53735 |
| BeTACoV/WuhAn/WIV05/2019\|EPI_ISL_402128 | 55171 | 55171 | 99% | 0.0 | 99.99% | Query_56711 |
| BeTACoV/WuhAn/WIV06/2019\|EPI_ISL_402129 | 55182 | 55182 | 99% | 0.0 | 100.00% | Query_60957 |
| BeTACoV/WuhAn/WIV07/2019\|EPI_ISL_402130 | 55171 | 55171 | 99% | 0.0 | 99.99% | Query_63943 |
| BeTACoV/WuhAn/HBCDC-HB-01/2019\|EPI_ISL_402132 | 55177 | 55177 | 99% | 0.0 | 100.00% | Query_50319 |
| IPBCAMS-WH-05/2020\|EPI_ISL_403928 | 55177 | 55177 | 99% | 0.0 | 100.00% | Query_53785 |
| BeTACoV/WuhAn/IPBCAMS-WH-04/2019\|EPI_ISL_403929 | 55182 | 55182 | 99% | 0.0 | 100.00% | Query_61767 |
| BeTACoV/WuhAn/IPBCAMS-WH-03/2019\|EPI_ISL_403930 | 55177 | 55177 | 99% | 0.0 | 100.00% | Query_64017 |
| BeTACoV/WuhAn/IPBCAMS-WH-02/2019\|EPI_ISL_403931 | 55149 | 55149 | 99% | 0.0 | 99.98% | Query_10691 |
| BeTACoV/GuAnGdonG/20SF012/2020\|EPI_ISL_403932 | 55166 | 55166 | 99% | 0.0 | 99.99% | Query_6757 |
| BeTACoV/GuAnGdonG/20SF013/2020\|EPI_ISL_403933 | 55166 | 55166 | 99% | 0.0 | 99.99% | Query_11145 |
| BeTACoV/GuAnGdonG/20SF014/2020\|EPI_ISL_403934 | 55177 | 55177 | 99% | 0.0 | 100.00% | Query_10319 |
| BeTACoV/GuAnGdonG/20SF025/2020\|EPI_ISL_403935 | 55166 | 55166 | 99% | 0.0 | 99.99% | Query_12643 |
| BeTACoV/GuAnGdonG/20SF028/2020\|EPI_ISL_403936 | 55177 | 55177 | 99% | 0.0 | 100.00% | Query_19609 |
| [BeTACoV/GuAnGdonG/20SF040/2020\|EPI_ISL_403937](https://blast.ncbi.nlm.nih.gov/Blast.cgi" \l "alnHdr_Query_25327" \o "Go to alignment for BeTACoV/GuAnGdonG/20SF040/2020\|EPI_ISL_403937) | 55177 | 55177 | 99% | 0.0 | 100.00% | Query_25327 |
| BeTACoV/NonThAburi/61/2020\|EPI_ISL_403962 | 55182 | 55182 | 99% | 0.0 | 100.00% | Query_40203 |
| BeTACoV/NonThAburi/74/2020\|EPI_ISL_403963 | 55182 | 55182 | 99% | 0.0 | 100.00% | Query_44483 |
| BeTACoV/ZhejiAnG/WZ-01/2020\|EPI_ISL_404227 | 55171 | 55171 | 99% | 0.0 | 99.99% | Query_46813 |
| BeTACoV/ZhejiAnG/WZ-02/2020\|EPI_ISL_404228 | 55182 | 55182 | 99% | 0.0 | 100.00% | Query_48655 |
| BeTACoV/USA/IL1/2020\|EPI_ISL_404253 | 55153 | 55153 | 99% | 0.0 | 99.97% | Query_6763 |
| BeTACoV/USA/WA1/2020\|EPI_ISL_404895 | 55166 | 55166 | 99% | 0.0 | 99.99% | Query_12809 |
| BeTACoV/Shenzhen/HKU-SZ-005/2020\|EPI_ISL_405839 | 55155 | 55155 | 99% | 0.0 | 99.98% | Query_53305 |
| BeTACoV/Shenzhen/HKU-SZ-002/2020\|EPI_ISL_406030 | 55166 | 55166 | 99% | 0.0 | 99.99% | Query_16689 |
| BeTACoV/TAiwAn/2/2020\|EPI_ISL_406031 | 55166 | 55166 | 99% | 0.0 | 99.99% | Query_56445 |
| BeTACoV/USA/CA1/2020\|EPI_ISL_406034 | 55143 | 55143 | 99% | 0.0 | 99.98% | Query_61555 |
| BeTACoV/USA/CA2/2020\|EPI_ISL_406036 | 55171 | 55171 | 99% | 0.0 | 99.99% | Query_44597 |
| BeTACoV/USA/AZ1/2020\|EPI_ISL_406223 | 55160 | 55160 | 99% | 0.0 | 99.99% | Query_47815 |

**Supplement Table 2: The model’s score by I-TASSER**

The confidence of each model is quantitatively measured by **C-score** that is calculated based on the significance of threading template alignments and the convergence parameters of the structure assembly simulations. **TM-score** and **RMSD** are estimated based on C-score and protein length following the correlation observed between these qualities.

| Name | **C-score** | **Exp.TM-Score e** | **Exp.RMSD** | **No.of decoys** | **Cluster density** |
| --- | --- | --- | --- | --- | --- |
| Model1 | -1.52 | 0.53+-0.15 | 13.3+-4.1 | 388 | 0.0271 |
| Model2 | -1.91 |  |  | 270 | 0.0184 |
| Model3 | -1.85 |  |  | 205 | 0.0194 |
| Model4 | -2.02 |  |  | 193 | 0.0165 |
| Model5 | -2.12 |  |  | 172 | 0.0149 |

**Supplement Table 3: Top 10 Identified stuctural analogs in PDB**

Ranking of proteins is based on TM-score of the structural alignment between the query structure and known structures in the PDB library. **RMSDa** is the RMSD between residues that are structurally aligned by TM-align. **IDENa** is the percentage sequence identity in the structurally aligned region. **Cov** represents the coverage of the alignment by TM-align and is equal to the number of structurally aligned residues divided by length of the query protein.

| **Rank** | **PDB-Hit** | **TM-Score** | **RMSDa** | **IDENa** | **CoV** |
| --- | --- | --- | --- | --- | --- |
| 1 | 5x58A | 0.825 | 0.76 | 0.733 | 0.0828 |
| 2 | 6nzkA | 0.743 | 4.60 | 0.269 | 0.0841 |
| 3 | 2vz9B | 0.278 | 10.11 | 0.034 | 0.0449 |
| 4 | 2pffD | 0.239 | 9.81 | 0.022 | 0.0374 |
| 5 | 3cu7A | 0.237 | 9.74 | 0.038 | 0.0369 |
| 6 | 2uv8G | 0.236 | 9.75 | 0.033 | 0.0368 |
| 7 | 1ea0A | 0.232 | 10.27 | 0.044 | 0.0375 |
| 8 | 2uvaG | 0.231 | 10.28 | 0.037 | 0.0374 |
| 9 | 1f31A | 0.227 | 9.94 | 0.025 | 0.0359 |

**Supplement Table 4: Top 5 pridection in Ligand bingding sites**

**C-score** is the confidence score of the prediction. **Cluster size** is the total number of templates in a cluster. **Lig Name** is name of possible binding ligand. **Rep** is a single complex structure with the most representative ligand in the cluster, i.e., the one listed in the Lig Name column. **Mult** is the complex structures with all potential binding ligands in the cluster

| **Rank** | **PDB-Hit Lig Name** | **C-scroe** | **Cluster size** | **Download Complex** | **Ligand Binding Site Residue** |
| --- | --- | --- | --- | --- | --- |
| 1 | 3srcA | 0.05 | 3 | 28N Rep,Mult | 874,877,878 |
| 2 | 1a5tA | 0.03 | 2 | ZN N/A | 738,743,749,760 |
| 3 | 4dv3B | 0.03 | 2 | MG Rep,Mult | 1004,1005 |
| 4 | 4fmaI | 0.03 | 2 | MG N/A | 907,1036,1037 |
| 5 | 5hhjA | 0.02 | 1 | GLY | 923,926 |

### Acknowledgment

We gratefully acknowledge the Originating and Submitting Laboratories for sharing newly identified coronavirus sequences through GISAID, as follows:

1 EPI_ISL_402119, EPI_ISL_402120, EPI_ISL_402121:

Originating and submitting lab - National Institute for Viral Disease Control and Prevention, China CDC

Authors - Wenjie Tan, Xiang Zhao, Wenling Wang, Xuejun Ma, Yongzhong Jiang, Roujian Lu, Ji Wang, Weimin Zhou, Peihua Niu, Peipei Liu, Faxian Zhan, Weifeng Shi, Baoying Huang, Jun Liu, Li Zhao, Yao Meng, Xiaozhou He, Fei Ye, Na Zhu, Yang Li, Jing Chen, Wenbo Xu, George F. Gao, Guizhen Wu

2 EPI_ISL_402123:

Originating and submitting lab - Institute of Pathogen Biology, Chinese Academy of Medical Sciences & Peking Union Medical College

Authors - Lili Ren, Jianwei Wang, Qi Jin, Zichun Xiang, Yongjun Li, Zhiqiang Wu, Chao Wu, Yiwei Liu

3 EPI_ISL_402124:

Originating lab - Wuhan Jinyintan Hospital Submitting lab - Wuhan Institute of Virology, Chinese Academy of Sciences

Authors - Peng Zhou, Xing-Lou Yang, Ding-Yu Zhang, Lei Zhang, Yan Zhu, Hao-Rui Si, Zhengli Shi

4 EPI_ISL_402125:

Originating lab - Unknown

Submitting lab - National Institute for Communicable Disease Control and Prevention (ICDC) Chinese Center for Disease Control and Prevention (China CDC)

Authors - Zhang,Y.-Z., Wu,F., Chen,Y.-M., Pei,Y.-Y., Xu,L., Wang,W., Zhao,S., Yu,B., Hu,Y., Tao,Z.-W., Song,Z.-G., Tian,J.-H., Zhang,Y.-L., Liu,Y., Zheng,J.-J., Dai,F.-H., Wang,Q.-M., She,J.-L. and Zhu,T.-Y.

5 EPI_ISL_402126:

Originating and submitting lab - Dept. of Virology III, National Institute of Infectious Diseases

Authors - Naganori Nao, Kazuya Shirato, Shutoku Matsuyama, Makoto Takeda

6 EPI_ISL_402127, EPI_ISL_402128, EPI_ISL_402129, EPI_ISL_402130, EPI_ISL_402132:

Originating lab - Wuhan Jinyintan Hospital

submitting lab - Hubei Provincial Center for Disease Control and Prevention

Authors - Lili Ren, Peng Zhou, Xing-Lou Yang, Ding-Yu Zhang, Lei Zhang, Yan Zhu, Hao-Rui Si, Zhengli Shi

6 EPI_ISL_403932, EPI_ISL_403933, EPI_ISL_403934, EPI_ISL_403935, EPI_ISL_403936, EPI_ISL_403937:

Originating lab – Guangdong Provincial Center for Diseases Control and Prevention; Guangdong Provincial Public Health

Submitting lab - Department of Microbiology, Guangdong Provincial Center for Diseases Control and Prevention

Authors - Min Kang, Jie Wu, Jing Lu, Tao Liu, Baisheng Li, Shujiang Mei, Feng Ruan, Lifeng Lin, Changwen Ke, Haojie Zhong, Yingtao Zhang, Lirong Zou, Xuguang Chen, Qi Zhu, Jianpeng Xiao, Jianxiang Geng, Zhe Liu, Jianxiong Hu, Weilin Zeng, Xing Li, Yuhuang Liao, Xiujuan Tang, Songjian Xiao, Ying Wang, Yingchao Song, Xue Zhuang, Lijun Liang, Guanhao He, Huihong Deng, Tie Song, Jianfeng He, Wenjun Ma

7 EPI_ISL_403962, EPI_ISL_403963:

Originating lab – Bamrasnaradura Hospital

Submitting lab - 1. Department of Medical Sciences, Ministry of Public Health, Thailand
2. Thai Red Cross Emerging Infectious Diseases - Health Science Centre
3. Department of Disease Control, Ministry of Public Health, Thailand

Authors - Pilailuk,Okada; Siripaporn,Phuygun; Thanutsapa,Thanadachakul; Supaporn,Wacharapluesadee; Sittiporn,Parnmen; Warawan,Wongboot; Sunthareeya,Waicharoen; Rome,Buathong; Malinee,Chittaganpitch; Nanthawan,Mekha

8 EPI_ISL_404227, EPI_ISL_404228:

Originating lab –Zhejiang Provincial Center for Disease Control and Prevention

Submitting lab - Department of Microbiology, Zhejiang Provincial Center for Disease Control and Prevention

Authors - Yin Chen, Yanjun Zhang, Haiyan Mao, Junhang Pan, Xiuyu Lou, Yiyu Lu, Juying Yan, Hanping Zhu, Jian Gao, Yan Feng, Yi Sun, Hao Yan, Zhen Li, Yisheng Sun, Liming Gong, Qiong Ge, Wen Shi, Xinying Wang, Wenwu Yao, Zhangnv Yang, Fang Xu, Chen Chen, Enfu Chen, Zhen Wang, Zhiping Chen, Jianmin Jiang, Chonggao Hu

9 EPI_ISL_404895:

Originating lab –Providence Regional Medical Center

Submitting lab - Division of Viral Diseases, Centers for Disease Control and Prevention

Authors - Queen,K., Tao,Y., Li,Y., Paden,C.R., Lu,X., Zhang,J., Gerber,S.I., Lindstrom,S., Tong,S.

10 EPI_ISL_405839,EPI_ISL_406030:

Originating lab –The University of Hong Kong - Shenzhen Hospital

Submitting lab - Li Ka Shing Faculty of Medicine, The University of Hong Kong

Authors - Chan,J.F.-W., Yuan,S., Kok,K.H., To,K.K.-W., Chu,H., Yang,J., Xing,F., Liu,J., Yip,C.C.-Y., Poon,R.W.-S., Tsai,H.W., Lo,S.K.-F., Chan,K.H., Poon,V.K.-M., Chan,W.M., Ip,J.D., Cai,J.P., Cheng,V.C.-C., Chen,H., Hui,C.K.-M. and Yuen,K.Y.

11 EPI_ISL_404253:

Originating lab –IL Department of Public Health Chicago Laboratory

Submitting lab - Pathogen Discovery, Respiratory Viruses Branch, Division of Viral Diseases, Centers for Dieases Control and Prevention

Authors - Ying Tao, Krista Queen, Clinton R. Paden, Jing Zhang, Yan Li, Anna Uehara, Xiaoyan Lu, Brian Lynch, Senthil Kumar K. Sakthivel, Brett L. Whitaker, Shifaq Kamili, Lijuan Wang, Janna' R. Murray, Susan I. Gerber, Stephen Lindstrom, Suxiang Tong

12 EPI_ISL_406031:

Originating and Submitting lab - Centers for Disease Control, R.O.C. (Taiwan)

Authors - Ji-Rong Yang, Yu-Chi Lin, Jung-Jung Mu, Ming-Tsan Liu, Shu-Ying Li

13 EPI_ISL_406034,EPI_ISL_406036:

Originating lab –California Department of Public Health

Submitting lab - Pathogen Discovery, Respiratory Viruses Branch, Division of Viral Diseases, Centers for Dieases Control and Prevention

Authors - Anna Uehara, Krista Queen, Ying Tao, Yan Li, Clinton R. Paden, Jing Zhang, Xiaoyan Lu, Brian Lynch, Senthil Kumar K. Sakthivel, Brett L. Whitaker, Shifaq Kamili, Lijuan Wang, Janna' R. Murray, Susan I. Gerber, Stephen Lindstrom, Suxiang Tong

13 EPI_ISL_406223:

Originating lab –Arizona Department of Health Services

Submitting lab - Pathogen Discovery, Respiratory Viruses Branch, Division of Viral Diseases, Centers for Dieases Control and Prevention

Authors - Ying Tao, Clinton R. Paden, Krista Queen, Anna Uehara, Yan Li, Jing Zhang, Xiaoyan Lu, Brian Lynch, Senthil Kumar K. Sakthivel, Brett L. Whitaker, Shifaq Kamili, Lijuan Wang, Janna' R. Murray, Susan I. Gerber, Stephen Lindstrom, Suxiang Tong
